# Digestion, fermentation, and pathogen anti-adhesive properties of the hMO-mimic di-fucosyl-β-cyclodextrin

**DOI:** 10.1101/2020.11.26.399972

**Authors:** Stella A. Verkhnyatskaya, Chunli Kong, Cynthia Klostermann, Henk A. Schols, Paul de Vos, Marthe T.C. Walvoort

## Abstract

**Scope:** Human milk is widely acknowledged as the best food for infants, and that is not just because of nutritional features. Human milk also contains a plethora of bioactive molecules, including a large set of human milk oligosaccharides (hMOs). Especially fucosylated hMOs have received attention for their anti-adhesive effects on pathogens by preventing attachment to the intestinal wall. Because hMOs are generally challenging to produce in sufficient quantities to study and ultimately apply in (medical) infant formula, hMO mimics are interesting compounds to produce and evaluate for their biological effects.

**Methods and results:** We investigated the digestion, fermentation, and pathogen anti-adhesive capacity of the novel hMO mimic di-fucosyl-β-cyclodextrin (DFβCD). We establish that DFβCD is not digested by α-amylase and also resists fermentation by the microbiota from a 9 month-old infant. In addition, we reveal that DFβCD blocks adhesion of enterotoxigenic *E. coli* (ETEC) to Caco-2 cells, especially when DFβCD is pre-incubated with ETEC prior to addition to the Caco-2 cells.

**Conclusion:** Our results suggests that DFβCD functions through a decoy effect. We expect that our results inspire the generation and biological evaluation of other fucosylated hMOs and mimics, to obtain a comprehensive overview of the anti-adhesive power of fucosylated glycans.

## 1. Introduction

Human milk is generally considered to be the best nutritional source for infants, as an increasing amount of research reveals the health effects of a plethora of bioactive components in human milk, including beneficial microbes,[1] immunoglobulins,[2] and human milk oligosaccharides (hMOs).[3, 4] Especially the hMOs have received considerable attention as health-promoting factors over the last decades, as they are shown to affect the development of the infant’s microbiome, promote immune system maturation, and ward off infections by acting as decoy substrates, amongst other effects.[5] Interestingly, hMOs are virtually absent from bovine milk-based infant formulas [6], and as a result, there is a broad interest to produce hMOs with the aim to add them to infant and medical formula. To achieve this goal, researchers are investigating the mode of action of specific hMOs to unravel structure-activity relationships that will guide the future selection of health-promoting hMO-based additives. Simultaneously, procedures are being developed to generate specific hMOs using (chemo)enzymatic and microbial cell factory approaches.[7] This has resulted in the recent approval by both the U.S. Food and Drug Administration (FDA) and the European Union of two short hMO structures, *i.e.* 2’-fucosyllactose (2’-FL) and lacto-*N*-neotetraose (LNnT), that are currently added to certain brands of infant formula.[8, 9]

Currently, more than 200 different hMO structures have been identified in human milk [10] and although they together form a complex mixture of carbohydrates, they do share certain structural similarities. HMOs are generally composed of a linear or branched backbone containing alternating *A*-acetyl-β-D-glucosamine (GlcNAc) and β-D-galactose (Gal) building blocks, capped with a lactose disaccharide on the reducing end. At specific positions these backbones can be decorated with α-L-fucose (Fuc) and α-D-neuraminic acid, also called sialic acid (Sia).[5] Comparing both types of decorations, it is interesting to note that Fuc is much more prevalent (on 50-80% of the hMOs) than Sia (on maximum 30% of hMOs).[11] Fucosylated hMOs have been linked to numerous health effects, including correcting microbial dysbiosis in cesarean-born infants[12] and protection against viral and bacterial infections[13] and bacterial gastroenteric infections.[14] Especially the anti-adhesive activity of fucosylated hMOs on pathogens is striking and reported by many.[15, 16] One of the first examples of the anti-adhesive potential of hMOs was published in 1990, where the neutral fraction (*i.e.* not containing Sia building blocks) of isolated hMOs showed a significant reduction in the adhesion of enteropathogenic *E. coli* (EPEC) and a concomitant preventative effect on the development of urinary tract infections.[17] In addition, maternal secretor status, which directly impacts the final structure and abundance of fucosylated oligosaccharide levels in the milk, was linked to infection rates and symptom severity in their offspring.[18] Using bioengineered samples of 2’-FL and 3-fucosyllactose (3-FL, both shown in **Figure 1**), moderate anti-adhesive effects against *Pseudomonas aeruginosa, Campylobacteri jejuni*, EPEC, and *Salmonella enterica* serovar *fyris* on intestinal Caco-2 cells were established.[19, 20] Moreover, 2’-FL was also found to block adhesion of *E. coli* O157 (an enterohemorrhagic *E. coli* strain, EHEC) onto intestinal Caco-2 cells.[21] Because the structural complexity of hMOs prevents their straightforward production in sufficient quantities to add to infant formula, structural analogs with similar functions have been developed, and with great success. Galacto-oligosaccharides (GOS) and fructo-oligosaccharides (FOS) are regularly added to infant formula, and induce especially the development of a healthy microbiome.[22, 23] Other so-called non-digestible carbohydrates (NDCs) include pectins, chitin and chitosan, alginates, and mannans, and many of these polysaccharides have been revealed to have anti-pathogenic effects. [24] As these NDCs are structurally highly different from native hMOs, this showcases that novel carbohydrate structures have the potential to mimic the beneficial effects of hMOs, without the challenging production that hMOs would require.

**Figure 1.**
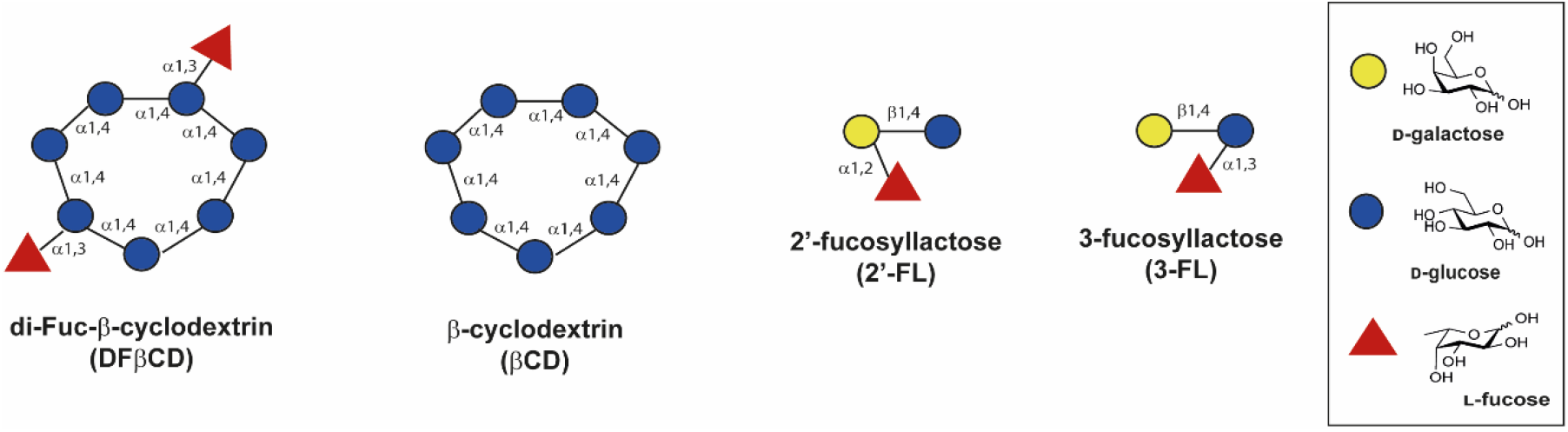
Overview of the molecules used in this study.

All this withstanding, the production of novel hMO analogs with enzymatic methods is currently an active area of research.[25] Enzymatic methods are currently mostly focused on hMOs containing either no or one decorative Fuc or Sia, including 2’-FL and 6’-sialyllactose (6’-SL).[26] With respect to fucosylated hMOs, details are emerging pertaining to the degree of backbone fucosylation and positioning on the backbone needed for a specific effect. For instance, 2’,3-di-fucosyllactose (DFL) exhibited a stronger antimicrobial effect against *Streptococcus agalactiae* GB590 than mono-fucosylated lactoses 2’-FL and 3-FL.[27] Especially a higher degree of fucosylation is nearly impossible to obtain with current enzymatic methods, compromising the ability to unravel the impact of higher degrees of fucosylation on anti-adhesion activity. Therefore, we recently developed a chemical strategy to produce di-fucosylated β-cyclodextrin (DFβCD, **Figure 1**).[28] The family of cyclodextrins has the GRAS status (Generally Recognized As Safe), and they are frequently used in medical formulations and as food additives.[29] β-Cyclodextrin (βCD) is shown in **Figure 1** and contains seven glucose (Glc) residues linked in an α-1,4 fashion, analogous to the structural composition of maltooligosaccharides. Using appropriately functionalized β-cyclodextrin and fucose building blocks, the di-fucosylated DFβCD was obtained in a highly regiospecific manner using chemical strategies.[28] Since the Fuc moieties are connected to the O-3 position of the backbone glucoses, the decorative pattern mimics the pattern in 3-FL (**Figure 1**), which also contains a Fuc moiety that is α-1,3 linked to glucose. As a result of this straightforward approach, DFβCD was produced in sufficient quantities (~ 0.5 g) to test its functional activity, and to determine whether this structural hMO mimic also is a functional hMO mimic.

Here we present our in-depth studies into the digestion and fermentation of DFβCD, as well as its anti-adhesive properties against enterotoxigenic *Escherichia coli* (ETEC) O78:H11. ETEC is the most common bacterial cause of diarrhea in children in developing countries, and although adhesion of ETEC to host cells is an intricate combination of factors,[30] it has been established that ETEC adhesion is also mediated by binding of bacterial lectins to host glycans.[31] In addition, 2’-FL was shown to reduce adhesion and invasion of ETEC bacteria to T84 intestinal epithelial cells *in vitro*.[32] Our results reveal that DFβCD is not digested and fermented by a 9 month-old infant’s inoculum, whereas it does have anti-adhesive properties against ETEC. This suggests that also hMO analogs such as DFβCD, composed of a different backbone structure but displaying appropriately spaced decorative Fuc moieties, can have similar health-beneficial effects as hMOs.

## 2. Experimental Section

### 2.1 Materials

The human milk oligosaccharides 2’-fucosyllactose (2’-FL) and 3-fucosyllactose (3-FL) were provided by Elicityl (France). Difucosylated β-cyclodextrin (DFβCD) was chemically synthesized based on the commercially obtained β-cyclodextrin (βCD), as described previously, and monofucosylated β-cyclodextrin (MFβCD) was isolated as a side product.[28] Pancreatin from porcine pancreas (containing amylase, lipase, and protease) and amyloglucosidase (260 U/mL) were obtained from Sigma-Aldrich (St. Louis, MO, USA). All materials needed for preparation of SIEM medium were obtained from Tritium Microbiology. *E. coli* ET8 was a gift from Prof. Gilles van Wezel (Leiden University)[33] and *E. coli* O78:H11 (ATCC35401) was purchased from ATCC. All chemicals used were of analytical grade.

### 2.2 *In vitro* digestibility of di-Fuc-β-cyclodextrin and β-cyclodextrin

Digestion was performed according to Martens et al. with minor modifications.[34] Di-Fuc-β-cyclodextrin (DFβCD) and β-cyclodextrin (βCD) were suspended in 100 mM sodium acetate buffer pH 5.9. Pancreatin solution was prepared according to Martens et al. [34], without the addition of invertase. In short, 150 mg pancreatin was suspended in 1 mL MQ and mixed for 10 min. The suspension was centrifuged for 10 min at 4 °C, 1500 x *g*. The final enzyme mixture was prepared by mixing 610 μL pancreatin supernatant with 58 μL amyloglucosidase and 83 μL MQ. Samples were incubated with 200 μL enzyme mixture for 0, 20, 60, 120, and 240 min at 20 mg/mL substrate concentration and enzymes were inactivated by boiling the sample for 15 min at 100 °C. Released glucose content was analysed with the GOPOD assay from Megazyme (Grey, Ireland).

### 2.3 *In vitro* fermentation of di-Fuc-β-cyclodextrin, β-cyclodextrin, 2’-FL and 3-FL

Fecal sample from one 9 month-old infant (vaginally born, breast-fed, introduced to solid food, no administration of antibiotics, exposed to probiotic Bifidobacteria) was collected and immediately stored in an anaerobic container. Fecal slurry was prepared by mixing fresh feces with a pre-reduced dialysate-glycerol solution according to Aguirre *et al.* [35] at 25 % w/v feces dialysate-10 % glycerol. The fecal slurry was snap-frozen in liquid nitrogen and stored at −80 °C. Standard Illeal Efflux Medium (SIEM) was prepared according to Logtenberg *et al.* [36] with minor modifications. The carbohydrate medium component contained (g/L): pectin, 12; xylan, 12; arabinogalactan, 12; amylopectin, 12; and starch, 100, with a final concentration of 0.592 g/L. The salt medium component contained 0.144 g/L NaCl. The inoculum was prepared by diluting the fecal slurry 25 times in SIEM medium. The substrate of interest was dissolved in SIEM medium at 2.22 mg/mL and 10 % inoculum was added. The *in vitro* fermentation was performed in the anaerobic chamber using sterile serum bottles. Samples were inoculated in duplicate and incubated for 0, 4, 8, 12, 24, and 36 h. In addition, blanks without substrate or without inoculum were incubated too. At each time point, 200 μL sample was taken from each bottle with a syringe. This sample was heated for 10 min at 100 °C to inactivate the enzymes and stored at −20 °C until analysis.

### 2.4 Substrate degradation analysis

Samples were diluted to 20 μg/mL (DFβCD and βCD) or 10 μg/mL (2’-FL and 3-FL) and centrifuged at 19000 x *g* for 10 min. The supernatant (10 μL injection volume) was analyzed using an ICS 3500 HPAEC system from Dionex (Sunnyvale, USA), in combination with a CarboPac PA-1 (2 x 250 mm) column, with a Carbopac PA-1 guard column (Dionex). Carbohydrate peaks were detected by an electrochemical Pulsed Amperometric detector (Dionex) after elution with 0.3 mL/min at 25 °C. The eluents consisted of A (0.1 M NaOH solution) and B (1 M NaOAc in 0.1 M NaOH). Two different gradients were used. For DFβCD and βCD the gradient used was: 2.5-25 % B (0-30 min), 25-100 % B (30-40 min), 100 % B (40-45 min), 2.5 % B (45-60 min). For 2’-FL and 3-FL the gradient used was 0-15 % B (0-15 min), 15-100 % B (15-20 min), 100 % B (20-25 min), 0 % B (25-45 min). Substrates were quantified using 2.5-10 μg/mL 2’-FL and 3-FL or 5-20 μg/mL DFβCD and βCD. In addition, mono-Fuc-β-CD (MFβCD) was injected as a pure compound to detect it within the DFβCD mix. Data analysis was performed with Chromeleon™ 7.2.6 software from Thermo Fisher Scientific (Waltham, Massachusetts, USA).

### 2.5 Organic acid formation *after in vitro* fermentation

Samples were diluted 5 times and centrifuged at 19000 x *g* for 10 min. The supernatant (10 μL injection volume) was analyzed using an Ultimate 3000 HPLC system from Dionex, in combination with an Animex HPX-87H column (Bio-Rad laboratories Inc, Hercules, USA). Samples were detected by a refractive index detector (RI-101, Shodex, Yokohama, Japan) and a UV detector set at 210 nm (Dionex Ultimate 3000 RS variable wavelength detector). Elution was performed at 0.5 mL/min and 50 °C using 50 mM sulphuric acid as eluent. Acetate, propionate, butyrate, lactate, and succinate standard curves were used for quantification (0.05 – 2 mg/mL). Data analysis was performed with ChromeleonTM 7.2.6 software from Thermo Fisher Scientific.

### 2.6 Culturing of the intestinal epithelial cell line

Human intestinal epithelial Caco-2 cells were cultured in Dulbecco’s Modified Eagle Medium (DMEM, Lonza), supplemented with 0.5% penicillin-streptomycin (50 μg/mL-50 μg/mL, Sigma), 1% non-essential amino acid (100x, Sigma), 10mM HEPES (Sigma), and 10% heat deactivated fetal calf serum (Invitrogen). Caco-2 cells between passage number of 15-20 were chosen for the experiment, and cells were routinely cultured in a humidified incubator with 5% CO_2_, at 37 °C. A number of 3 ×10^4^ Caco-2 cells were seeded onto 24-well plates and cultured for 21 days before use.

### 2.7 Culturing of bacterial cells

Pathogenic bacteria of *Escherichia (E.) coli* ET8, and *E. coli* O78: H11 were recovered from glycerol stocks at −80 °C overnight, and were cultured in Brain heart infusion (BHI). After recovery culture, *E. coli* ET8 and *E. coli* O78: H11 were plated on BHI agar. A single colony of each bacterium was inoculated from the agar plates to BHI broth for a second overnight culture at 37 °C before the adhesion assay.

### 2.8 Bacterial adhesion assay

All compounds were dissolved into 2, 5, and 10 mg/mL in antibiotics-free cell culture medium with 1% dimethyl sulfoxide. The cell culture medium without the tested molecules served as control. All compounds were heated for 30 min at 65 *°C* to remove any endotoxin contamination before use. We first tested the concentration-dependent effect of the molecules on the adhesion of one pathogen *E. coli* ET8 to Caco-2 cells. This was done by pre-incubation of Caco-2 cells with 2’-FL, 3-FL, βCD, and DFβCD of 2, 5, and 10 mg/mL for 2h, respectively. *E. coli* ET8 was collected after 2h of culture with centrifugation at 2000 x *g* for 10 min and washed one time with PBS. The optical density (OD) of *E. coli* ET8 was adjusted to OD_540_ = 0.6 in PBS, and re-suspended either with or without the molecules at different concentrations. After that Caco-2 cells were co-incubated with the pathogens for another 2h at 37 °C. Afterwards, the non-adherent bacteria were washed away for three times in PBS. The adherent bacteria were released in 200 μL of 0.1% Triton-X100, and underwent serial dilutions in PBS. The drop-plating method [37] was employed to plate the adherent bacteria on BHI agar plates (n = 3). Then the molecules were applied at a concentration of 10 mg/mL to test the influence of the molecules on the adhesion of *E. coli* O78: H11 to Caco-2 cells. Here, to test the possible effects of the molecules on the adhesion, we either pre-incubated the molecules with Caco-2 cells to explore whether they could influence the bacteria adhesion through modification of the receptors on gut epithelial cells, or we pre-incubated the molecules with the bacteria to determine a possible decoy effect on the bacteria.

#### Pre-incubate with Caco-2 cells

As performed with *E. coli* ET8, all molecules at 10 mg/mL were pre-incubated with Caco-2 cells for 24h. *E. coli* O78: H11 was collected after 2h of subculture, when a log phase was reached, by centrifugation at 2000 x *g* for 10 min and the cells were washed with PBS. The OD of *E. coli* O78: H11 was adjusted to OD_540_ = 0.6 in PBS, and re-suspended either with or without the molecules and infected Caco-2 cells for another 2h at 37 °C.

#### Pre-incubate with bacteria. E. coli O78

H11 was collected as described above, pre-incubated with the molecules for 2h at 37 *°C* after re-suspension, and then infected Caco-2 cells for another 2h at 37 °C.

After infection, Caco-2 cells were gently washed three times to remove the non-adherent bacteria, and the adherent bacteria were collected with 200 μL of 0.1% Triton-X100, followed by serial dilutions in PBS. Total colony-forming units (CFUs) were determined after drop-plating method (n = 5).

### 2.9 Statistical analysis

Statistical analysis was performed with GraphPad Prism 6 (GraphPad Prism LLC.). Normality of the data distribution was confirmed with Kolmogorov-Smirnov test. The results were expressed as mean ± SD. All data were analyzed with Kruskal-Wallis test of one-way ANOVA, except for the decoy effect test of *E. coli* O78: H11, which was done with RM one-way ANOVA. Significant difference was defined as p<0.05 (*p<0.05), p<0.1 was considered as a statistical trend.

## 3. Results

### 3.1 HPAEC analysis of the hMOs and DFβCD

We first analyzed the chromatographic behavior of the compounds under study here. All compounds were subjected to High-Performance Anion-Exchange Chromatography (HPAEC) coupled to a Pulsed Amperometric Detector (PAD), and an overview of the respective masses and retention times (*R*t) is given in **Table 1** and the chromatograms are shown in **Figure S1**. 2’-FL and 3-FL were analyzed using gradient A, while βCD and difucosylated βCD (DFβCD) were analyzed using gradient B. Owing to the different masses and interactions with the column, the retention times on the HPAEC column differed greatly. The DFβCD sample was shown to contain 20% of monofucosylated βCD (MFβCD), which was confirmed by comparing the retention time with that of a pure MFβCD sample. The HPAEC elution pattern showed that DFβCD (16.7 min) was accompanied by a minor peak at 15.7 min (Figure S1), presumably corresponding to an isomer with a different fucose substitution pattern.

**Table 1.** Retention times of the hMOs and hMO mimics used in this study.

| | Compound | Exact mass (Da) | $R_t$ (min) |
| --- | --- | --- | --- |
| Gradient A | 2'-FL | 488.1741 | 8.2 |
|  | 3-FL | 488.1741 | 5.7 |
| Gradient B | βCD | 1134.3698 | 29.2 |
| | MFβCD | 1280.4277 | 21.0 |
| | DFβCD | 1426.4856 | 15.7, 16.7 |

Gradient A: 0-15 % B (0-15 min), 15-100 % B (15-20 min), 100 % B (20-25 min), 0 % B (25-45 min);
Gradient B: 2.5-25 % B (0-30 min), 25-100 % B (30-40 min), 100 % B (40-45 min), 2.5 % B (45-60 min)

### 3.2 DFβCD is resistant to digestion by pancreatic enzymes

Human milk oligosaccharides are generally characterized as non-digestible carbohydrates because they resist digestion by salivary α-amylase and digestive enzymes of the small intestine.[38, 39] Consequently, a prerequisite for novel hMO-type compounds to exert beneficial effects in the intestinal environment is that they also resist enzymatic digestion in the upper part of the gastrointestinal tract. [38] Because DFβCD is a starch-like compound composed of α-1,4-linked glucose units (**Figure 1**), it may be digested by salivary α-amylase or pancreatic α-amylase in the small intestine. To determine the resistance to digestion, DFβCD and β-cyclodextrin (βCD) were digested in an *in vitro* model according to Martens *et al.* [40] and the amount of released glucose was quantified over time (**Figure 2**).

**Figure 2.**
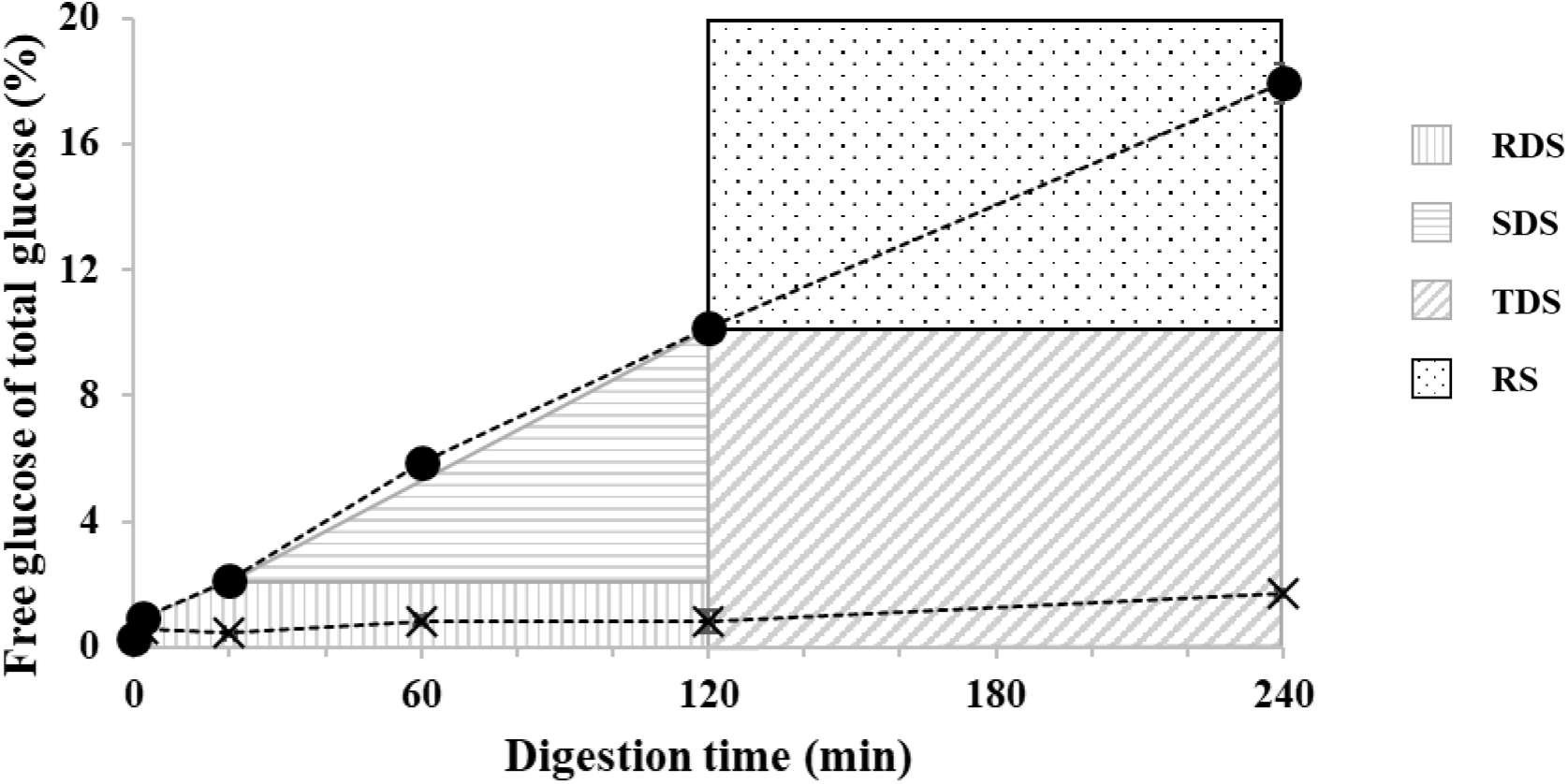
*In vitro* digestibility of DFβCD (×) and βCD (•) during 240 min of incubation expressed as free (released) glucose of total glucose (%). Soluble potato starch was digested for 90% within 120 min of incubation (Figure S2). For reference, rapidly digestible starch (RDS), slowly digestible starch (SDS), total digestible starch (TDS), and resistant starch (RS) are indicated in the figure.

The results show that DFβCD is almost fully resistant to digestion by pancreatic enzymes. Interestingly, βCD was also quite resistant towards digestion, as only 10% digestion was observed within 120 min of incubation, whereas the other 90% can be recognized as resistant starch according to the definition of Englyst *et al*.[41] Apparently, the cyclic form has a large influence on the binding of the substrate by digestive enzymes. For comparison, soluble potato starch was digested as a positive control, which was 90% hydrolyzed within 120 min of incubation (**Figure S2**). From the curve in **Figure 2**, it can be expected that βCD would be fully degradable by pancreatic enzymes upon extended incubation times.

### 3.3 DFβCD is resistant to bacterial fermentation and does not induce short-chain fatty acid production

Having established that DFβCD resists enzymatic digestion and is likely to arrive in the colon intact, we subsequently tested the possible fermentability of DFβCD by infant microbiota. All compounds were initially subjected to a fermentation experiment *in vitro* for 36 h using the fecal inoculum of a 9 month-old infant, and the extent of fermentation over time was quantified by the disappearance of the compound peak using HPAEC-PAD analysis (**Figure 3A**).

**Figure 3.**
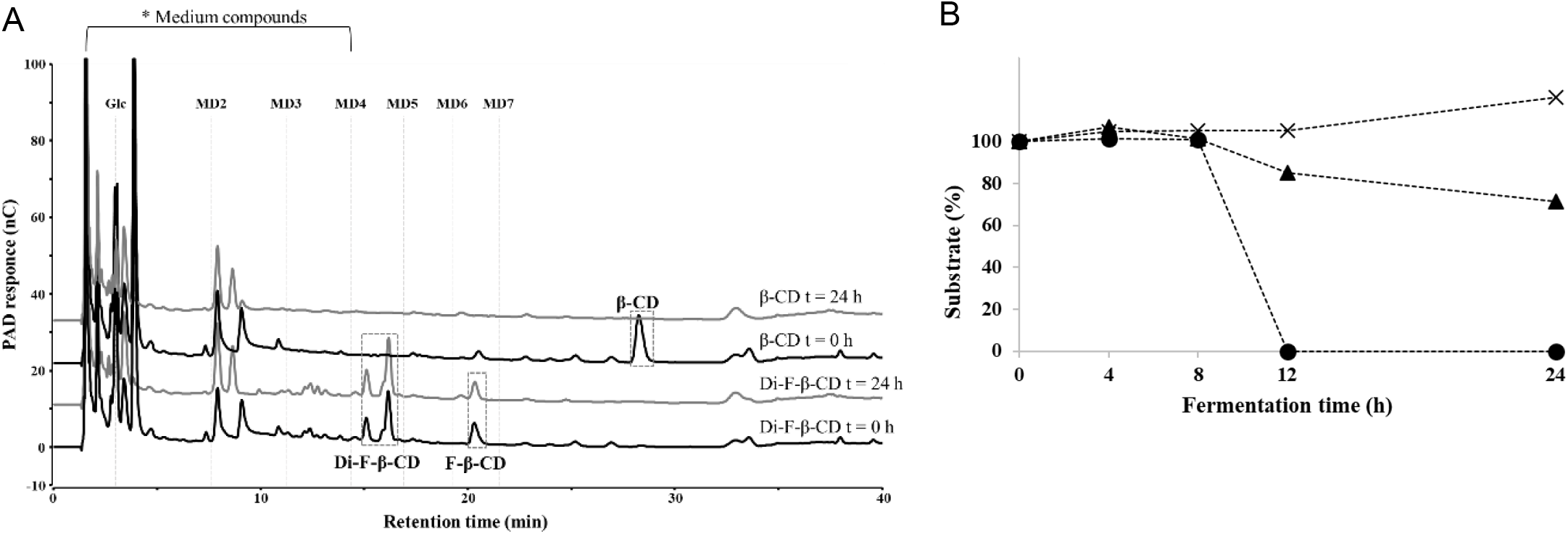
Fermentation studies. A) HPAEC elution pattern of DFβCD and βCD at 0 and 24 h of *in vitro* fermentation by 9-month old infant inoculum. Medium compounds are indicated with an *, malto-oligomers are indicated with Glc, and MD2-7. B) Time-dependent *in vitro* fermentation of DFβCD (╳), MFβCD (▲) and βCD (•) during 24 h.

Interestingly, whereas unsubstituted βCD was completely utilized after 24 h of fermentation, no degredation was observed for DFβCD. A small decrease in the peak area of MFβCD was observed between t = 0 h and t = 24 h, suggesting the fermentation was very slow (**Figure 3A**). As a comparison, the fermentation experiment was also conducted on 2’-FL and 3-FL, and both hMOs were completely degraded after 24 h of fermentation (**Figure S3**). Substrate degradation was monitored closely over time to assess the fermentation kinetics (**Figure 3B**). DFβCD was indeed not degraded at all by microbial enzymes of a 9-month old infant’s microbiome during batch fermentation. However, it seems that the degradation of mono-fucosylated MFβCD, which is present for 20 % within the DFβCD mix, started between 8-12 h of incubation reaching ±70% fermentation after 24 h. Degradation of βCD started after 8 h of incubation quite suddenly and was completed at 12 h of incubation. No decrease in substrate degradation was observed until 8 h of incubation and also no intermediate malto-oligomers were detected by HPAEC-PAD at any time of incubation. This delay may suggest that opening of the CD ring is the limiting step for successful degradation. 2’-FL and 3-FL were both degraded by the microbiota of this 9-month old infant (**Figure S3B**), and the degradation of 2’-FL seemed to be slightly faster than that of 3-FL.

After fermentation of the four compounds using a 9-month old infant inoculum, the production of short-chain fatty acids (SCFA) was analyzed using HPLC-RI-UV (**Figure S4**). In the case of DFβCD, little change in SCFA production compared to the fermentation control in medium was observed (**Figures S4A** and **S4E**). This was as expected based on the observed lack of degradation of DFβCD (**Figure 3**). Fermentation of unsubstituted βCD resulted in faster SCFA production compared to the fermentation control, and a higher amount of butyrate was produced (acetate:butyrate 67:32 compared to 80:20 at 12 h, **Figures S4B** and **S4E**). Fermentation of the human milk oligosaccharides 2’-FL and 3-FL also resulted in a quite fast production of acetate and intermediate acids lactate and succinate (for 2’-FL the ratio is acetate:butyrate:lactate:succinate 65:13:13:8, and for 3-FL the ratio is 67:11:14:7 at 12 h, **Figures S4C** and **S4D**), which were further converted to acetate and butyrate over the course of the experiments.

### 3.4 DFβCD reveals structure-dependent anti-adhesive effects against *E. coli* O78:H11

To investigate the anti-adhesive properties of DFβCD in comparison with 2’-FL, 3-FL, first an adhesion assay with the laboratory *E. coli* strain ET8 was performed. In this experiment, Caco-2 cells were pre-incubated with the compounds at concentrations of 2, 5, and 10 mg/mL, followed by exposure to *E. coli* ET8 bacteria and quantification of adhered bacteria (**Figure S5**). Interestingly, of all compounds tested only DFβCD at 10 mg/mL significantly inhibited the adhesion of *E. coli* ET8 to intestinal epithelial Caco-2 cells, with a reduction in adhesion of 60% compared to the non-treated control (Figure S5C, p<0.05). Subsequently, ETEC O78:H11 (H10407) was selected as a clinically relevant pathogen to assess the anti-adhesive capacity of DFβCD. We started by pre-incubating Caco-2 cells with 2’-FL, 3-FL, βCD and DFβCD for 24 h, followed by the addition of ETEC bacteria (log phase). Bacterial adhesion in the presence of DFβCD and control hMOs was determined by counting the CFU per mL of serially diluted washes from Caco-2 monolayers, as compared to the control incubation (no additive). Using the control reaction, the fraction of adhering bacteria was corrected for continuous bacterial growth in the assay. As shown in **Figure 4A**, when the Caco-2 cells were pre-incubated with the molecules for 24 h before *E. coli* O78:H11 infection, we observed that 2’-FL, 3-FL, βCD and DFβCD all reduced the adhesion of *E. coli* O78:H11 to Caco-2 cells. The decrease in inhibition was calculated to be 37%, 43%, 34%, 16%, respectively, of which only the adhesion reduction of 3-FL achieved statistical significance (p < 0.05). Both 2’-FL and βCD showed a trend of inhibition of adhesion (p values of 0.06 and 0.09, respectively).

In contrast, when the ETEC bacteria were first pre-incubated with the compounds and subsequently added to confluent Caco-2 cells, 2’-FL, 3-FL, and DFβCD all significantly inhibited the adhesion of *E. coli* O78:H11 to Caco-2 cells, with a reduction of 30% (p < 0.05), 21% (p < 0.05), and 42% (p < 0.05), respectively (**Figure 4B**). Of all compounds tested here, DFβCD actually revealed the highest inhibition of adhesion. These results are fucose-dependent, as the non-fucosylated βCD did not significantly impact bacterial adhesion.

**Figure 4.**
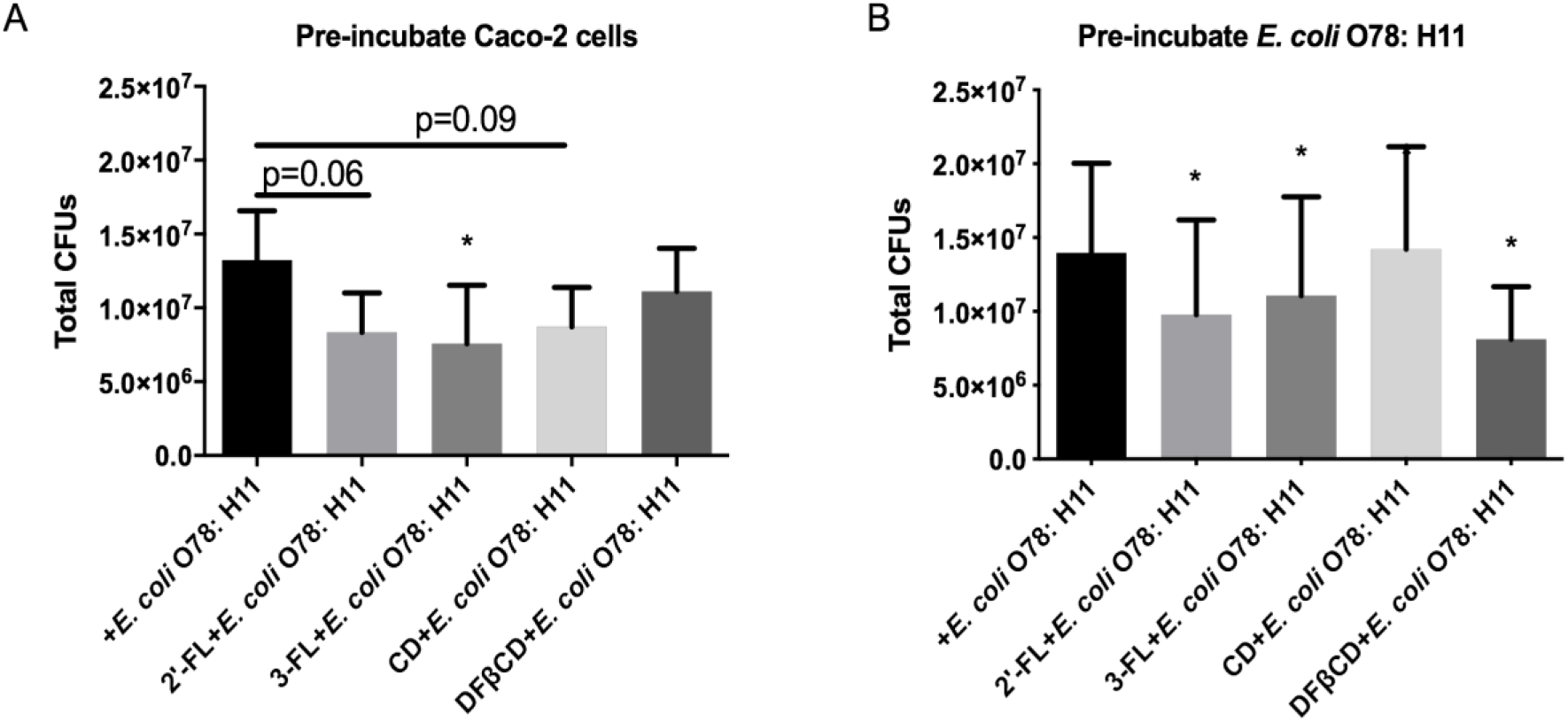
hMOs and DFβCD in 10 mg/mL inhibited adhesion of *E. coli* O78: H11 to intestinal epithelial Caco-2 cells. Caco-2 cells were cultured in 24 well plates for 21 days. The tested molecules of 2’-FL, 3-FL, βCD, and DFβCD either pre-incubated with Caco-2 cells for 24h (A) or pre-incubated with *E. coli* O78: H11 for 2h (B). Cell culture medium without tested molecules was taken as control.

## 4. Discussion

The increasing evidence that human milk oligosaccharides inhibit bacterial infections has led to a surge in interest to understand the exact structure-activity relationships of specific hMOs, and to develop methods to generate these structures. Especially the fucosylated hMOs have been linked to anti-pathogenic effects,[42, 43] and this may be a result of the central role of fucose on extracellular glycans that are involved in a plethora of biological functions, including serving as anti-adhesion molecules for pathogens.[44] As hMO structures in general, and multiply fucosylated hMOs specifically, are challenging to produce on large scale and with high purity, there is a high demand for alternative compounds that elicit similar effects.[45]

Here we report the biological evaluation of the novel hMO mimic di-fucosyl-β-cyclodextrin (DFβCD). Using enzymatic digestion and *in vitro* fermentation analyses, we established that DFβCD is resistant to digestive enzymes, and as a result is expected to reach the large intestine intact. Apparently, the digestive enzymes, that generally hydrolyze α-1,4-linked Glc units that are also present in DFβCD, are blocked from action by the two Fuc units. As non-fucosylated βCD is only slowly digested, the cyclic structure of βCD is also hypothesized to have a major impact on the resistance. This fits well with earlier reports that βCD is only slowly digested by α-amylase, while αCD (six glucose units) is resistant and γCD (8 glucose units) is quickly digested.[46] Both 2’-FL, 3-FL, and βCD are efficiently fermented by microbial enzymes, whereas DFβCD is resistant to bacterial fermentation. We hypothesize that the resistance observed for DFβCD is a result of the specific positioning of the two Fuc units on opposite sides of the βCD ring (specifically at the A and D position). [28] This may block the accessibility to the microbial α-amylases sufficiently to resist fermentation, at least over the course of the 24 h incubation. Overall SCFA production in all fermentated substrates after 24 h did not differentiate much from the levels of SCFAs produced in the medium-only samples, presumably due to the low substrate concentration used (2 mg/mL). [36] Slight differences in level and relative concentrations of the various acids at 12 h of fermentation can be seen, and are substrate dependent. Fermentation experiments at higher concentrations (*e.g.*, 10 mg/mL) may provide a more accurate picture of the levels of SCFAs produced with 2’-FL, 3-FL, and βCD over time. From the resistance to digestion observed with DFβCD it is expected that for this compound the absence of additional SCFA production will be confirmed.

Excitingly, DFβCD revealed anti-adhesive properties against enterotoxigenic *E. coli* strain O78:H11. There are two major mechanisms that may be at the basis of the anti-adhesive effect of DFβCD, *i.e.* through modulating intestinal cell susceptibility to bacterial adhesion by changing receptor expression levels, or through direct scavenging of bacteria by serving as decoy substrates. Both possible mechanisms were investigated by pre-incubating the compounds with the Caco-2 cells or the bacterial cells, respectively. Because the anti-adhesive effect of DFβCD was especially apparent when pre-incubated with the bacteria prior to exposure to Caco-2 intestinal epithelial cells, we postulate that DFβCD acts as a decoy substrate. Upon pre-incubation of *E. coli* O78:H11 with DFβCD, the hMO mimic may saturate the receptors on the bacterial cell surface, and thereby prevent binding to the glycans on Caco-2 intestinal cells. Such binding would require the pathogen to express specific receptors that generally bind epithelial cell surface-associated glycans, but when confronted with DFβCD, may bind this compound instead. When the Caco-2 intestinal cells were first pre-incubated with DFβCD, a less pronounced effect was observed, suggesting that DFβCD has little impact on the expression of cell-surface proteins involved in adhesion.

One of the first steps in pathogen colonization is adhesion to host cells, and its ability to adhere is directly correlated with the pathogen’s capacity to invade and infect. Fucose-dependent pathogens have been shown to be prevented by fucose-containing hMOs from adhering to mucosal membranes.[47] Interfering with pathogen adhesion is therefore an effective strategy to prevent infection.[48] Whereas natural hMOs are ideal candidates for anti-adhesive compounds, hMO mimics also have a large potential to fill the current void between functional relevance and availability of the natural hMO compounds. The biological activity of DFβCD serves as a strong proof-of-concept that hMO mimics are capable of eliciting anti-adhesive effects similar to hMOs. The generation and evaluation of other fucosylated structures will further our knowledge on the potential of fucosylated compounds to block bacterial infections.

## Supporting information

Supp Info Walvoort_2FucCD_manuscript

## Supporting Information

Supplementary figures are available in a separate file.

## Author contributions

M.T.C.W., S.V., P.d.V. and H.A.S. conceived the project. S.V. generated the DFβCD compound. C.Ko. performed the anti-adhesion studies. C.Kl. performed the digestion and fermentation studies. C.Ko., C.Kl, P.d.V., H.A.S., and M.T.C.W. were involved in data analysis. M.T.C.W. and S.V. wrote the manuscript with input from all authors. All authors were involved in proofreading the manuscript.

## Acknowledgments

This work was financially supported by the European Union through the Rosalind Franklin Fellowship COFUND project 60021 (to M.T.C.W.), and by the China Scholarship Council (CSC) grant number 201600090212 (to C.Ko).

## List of Abbreviations

βCD: β-cyclodextrin
DFβCD: di-Fuc-β-cyclodextrin
DFL: 2’,3-difucosyllactose
EHEC: enterohemorrhagic *E. coli*
EPEC: enteropathogenic *E. coli*
ETEC: enterotoxigenic *E. coli*
2’-FL: 2’-fucosyllactose
3-FL: 3-fucosyllactose
FOS: fructo-oligosaccharides
Fuc: L-fucose
Gal: D-galactose
Glc: D-glucose
GlcNAc: *A*-acetyl-D-glucosamine
GOS: galacto-oligosaccharides
hMO: human milk oligosaccharide
MFβCD: mono-Fuc-β-cyclodextrin
NDC: non-digestible carbohydrate
Sia: D-sialic acid
6’-SL: 6’-sialyllactose

