## Supplementary material for "Digestion, fermentation, and pathogen anti-adhesive properties of the hMO-mimic di-fucosyl-β-cyclodextrin": Supp Info Walvoort_2FucCD_manuscript

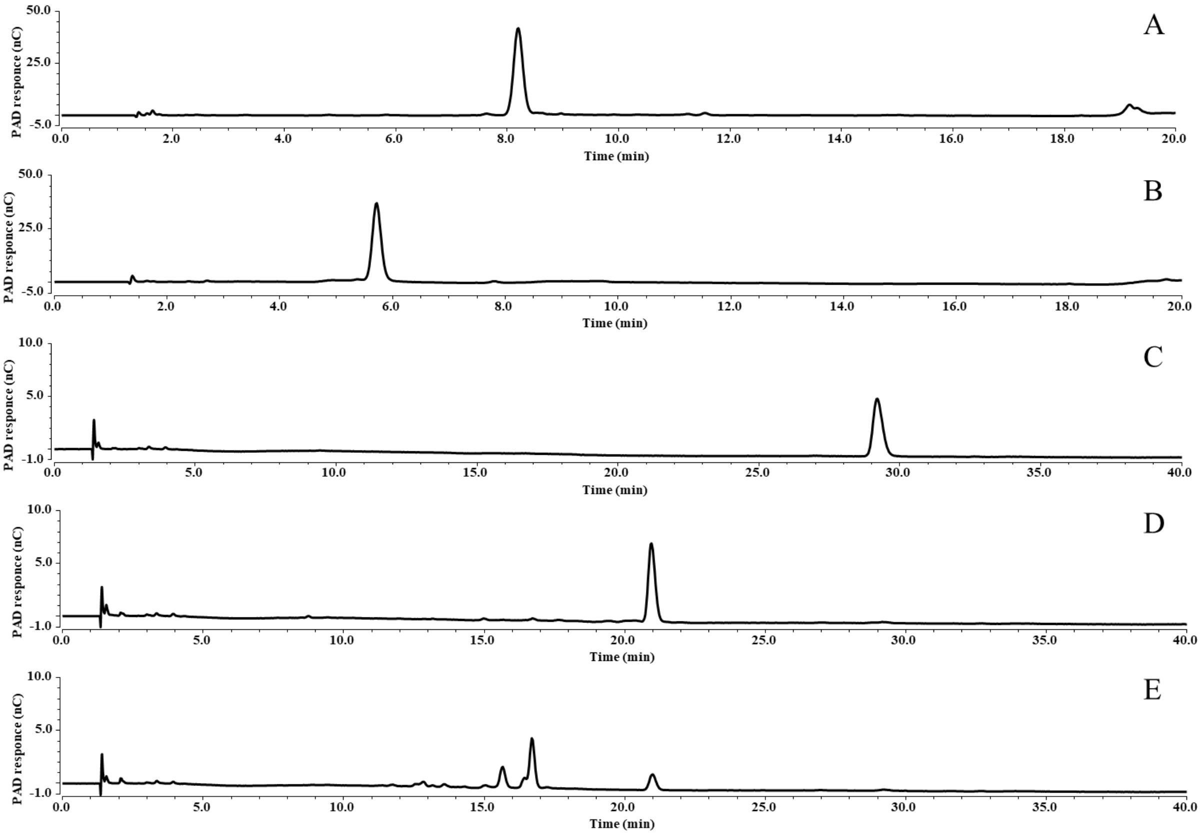


**Figure S1**. **HPAEC chromatography traces of the compounds used in this study.** A, B, C, D and E refer to 2’-FL, 3-FL, βCD, MFβCD, and DFβ-CD, respectively. Different gradients were used for A-B and C-E, see materials and methods section.

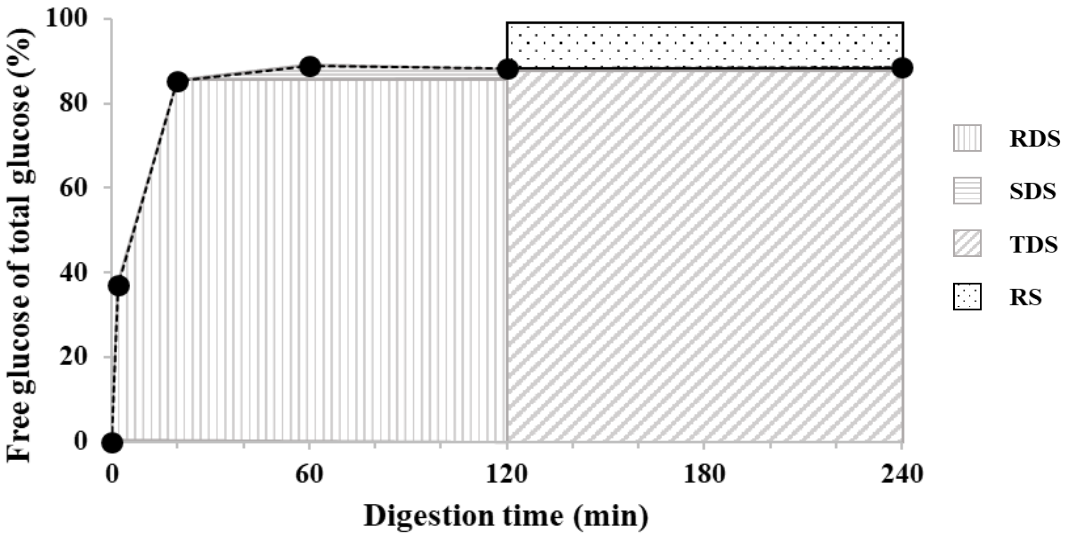


**Figure S2.** *In vitro* digestibility of soluble potato starch during 240 min of incubation expressed as free (released) glucose of total glucose (%).

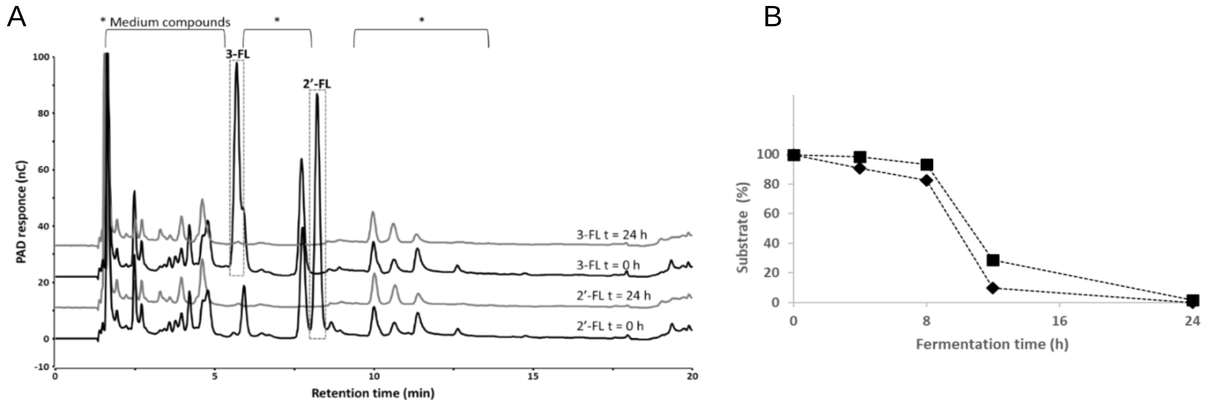


**Figure S3. Fermentation studies of 2'-FL and 3-FL.** A) HPAEC elution pattern of 2'-FL and 3-FL at 0 and 24 h of *in vitro* fermentation by 9-month old infant inoculum. Medium compounds are indicated with an *. B) Time-dependent *in vitro* fermentation of 2'-FL (◆) and 3-FL (■) during 24 h.

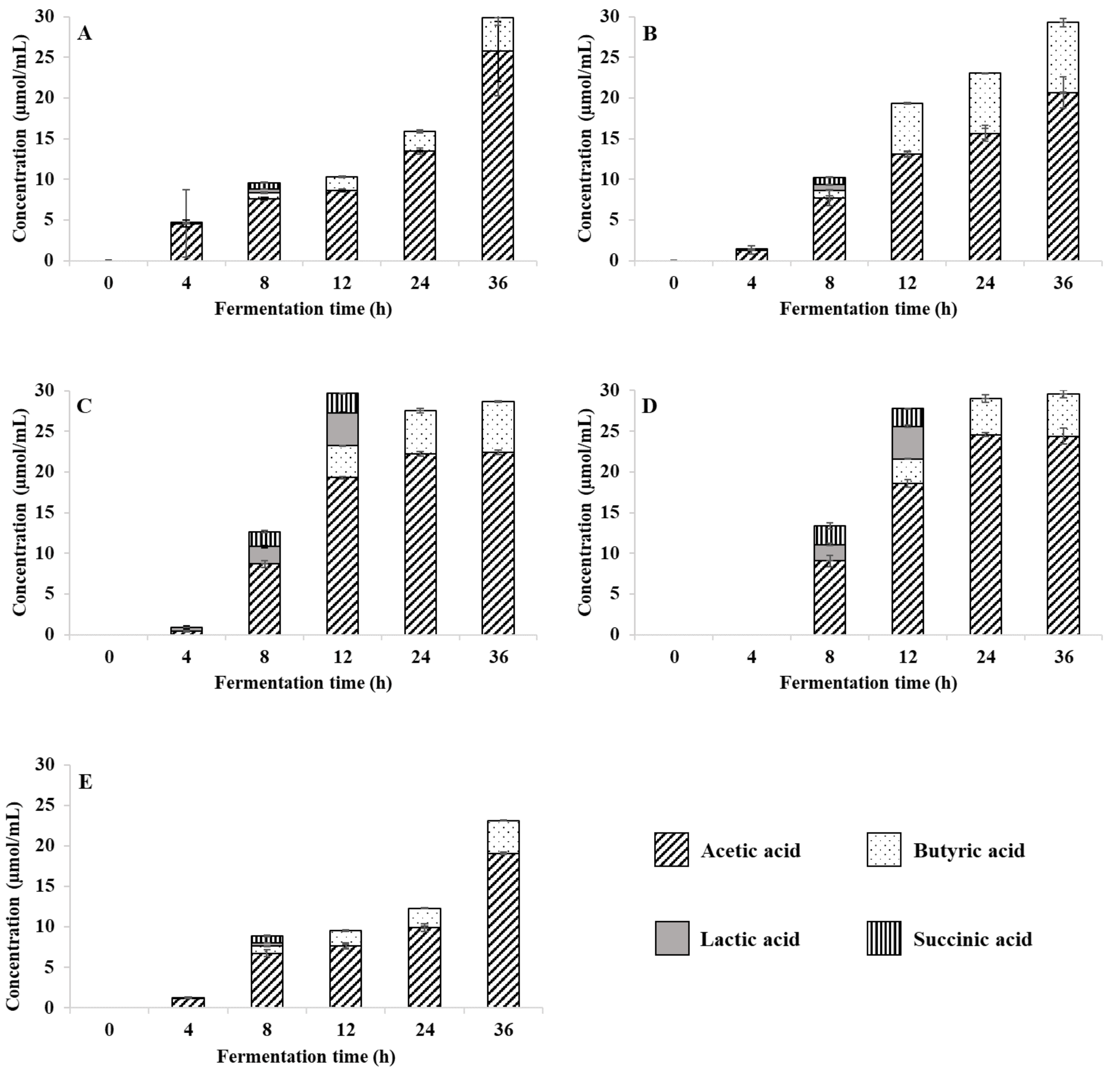


**Figure S1. Short-chain fatty acid formation (µmol/mL fermentation medium) during *in vitro* fermentation using 9 month-old infant inoculum.** A) DFβCD including 20 % MFβCD, B) βCD, C) 2’-FL, D) 3-FL and E) SIEM medium without substrate.

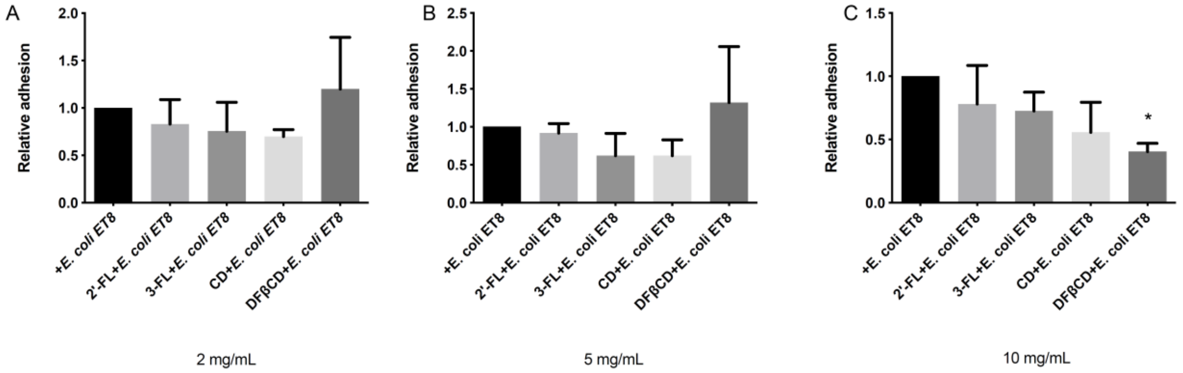


**Figure S5. DFβCD inhibited adhesion of *E. coli* ET8 to intestinal epithelial Caco-2 cells in a concentration-dependent manner**. Caco-2 cells were cultured in 24 well plates for 21 days and pre-incubated with 2'-FL, 3-FL, βCD, and DFβCD at 2 (A), 5 (B), 10 (C) mg/mL for 2h before infection of *E. coli* ET8. Cell culture medium without tested molecules was taken as control. After another 2h of infection, the total colony-forming units (CFUs) adhered to Caco-2 cells were determined by the drop-plating method. All data were normalized and were expressed as mean ± SD from three experiments. Statistical significance was tested with one-way ANOVA (*$p < 0.05$).
